# p53 and RB Cooperate to Suppress Transposable Elements

**DOI:** 10.1101/2023.02.06.527304

**Authors:** Omar M. Lopez, Michelle Zhang, Peter G. Hendrickson, Jianguo Huang, Andrea R. Daniel, Jane Blackmer, Lixia Luo, Laura D. Attardi, David Corcoran, David G. Kirsch

**Affiliations:** Department of Pharmacology and Cancer Biology, Duke University School of Medicine, Durham NC; Department of Radiation Oncology, Duke University School of Medicine, Durham NC; Department of Genetics, University of North Carolina at Chapel Hill, Chapel Hill NC; Division of Radiation and Cancer Biology, Department of Radiation Oncology, Stanford University School of Medicine, Stanford, CA; Department of Genetics, Stanford University School of Medicine, Stanford

## Abstract

Accumulating data suggest that the long interspersed nuclear element 1 (LINE1), a major class of transposable elements, can promote tumorigenesis by causing deletions of tumor suppressor genes, amplifications of oncogenes, and chromosomal translocations(1). LINE1 and other transposable elements such as short interspersed nuclear elements (SINEs) can be activated during cancer development. Aberrant transposon expression has been correlated with loss of p53, however, how the loss of p53 or other tumor suppressors leads to activation of transposons during tumorigenesis remains to be fully understood. Here, we investigate the link between loss of the p53 and RB tumor suppressors and derepression of transposable elements. We observe that loss of p53 and RB in mouse embryonic fibroblasts leads to epigenetic changes, such as loss of DNA and histone methylation, which correlate with marked upregulation of LINE and SINE transposable elements. These results suggest that p53 and RB may utilize epigenetic mechanisms that work in tandem to suppress transposons, which may contribute to tumor suppression.

**Significance:** The two most commonly mutated genes in human cancer are *p53* and the retinoblastoma (*RB*) tumor suppressors. Together, these genes regulate complex interconnecting pathways responsible for the regulation of cell growth, cell death, and genomic integrity. Studies in genetically engineered mouse models demonstrate that the canonical functions of these two tumor suppressors fail to fully explain their tumor suppressive capabilities. Individually, p53 and RB have each been implicated in repressing transposons. Here we show that combined loss of p53 and RB, which rapidly transforms cells, also results in cooperative upregulation of transposable elements and coincides with decreased methylation of DNA and H3K9. Our results present a framework by which p53 and RB work together to restrain the expression of LINE1 and SINEs, which may contribute to tumor suppression.

## Introduction

Repetitive elements make up over 50% of the human genome and the majority of these are transposable elements (TEs) (2) (Fig. 1A). TEs fall into two dichotomous classes:retrotransposons (RTs), which mobilize via a copy-and-paste mechanism, or DNA transposons, which propagate via a cut-and-paste method. The LINE1 family belongs in the former class and comprises a large fraction (~17%) of human genomic DNA. During embryogenesis, LINE1 expression and retrotransposition occurs as a consequence of the extensive chromatin remodeling process that is required for gametic reprogramming (3).Thereafter, somatic cells maintain several redundant LINE1 silencing mechanisms, which include histone methylation, DNA methylation and RNA silencing (4, 5). The best studied mechanism of TE repression involves over 200 KRAB-Zinc Finger Proteins (KRAB-ZFPs) that can specifically recognize TEs and recruit a series of epigenetic modifiers including DNA methyltransferases (DNMT3A/B) to methylate DNA as well as histone methyltransferases (SUV39h1) that deposit, among other repressive marks, H3K9me3 (4–6). Additionally, a Rb-EZH2 complex has been shown to mediate TE silencing by directing H3K27me3 deposition via the polycomb repressor 2 complex (PRC2) (7, 8). As an added layer of defense, mammals encode four RNA binding PIWI proteins which together with their RNA partners (piRNAs) target TE cDNA for degradation(6). piRNAs are expressed mainly in the germline of spermatogenic cells, but recent studies have identified them in neurons as well (9).

**Figure 1.**
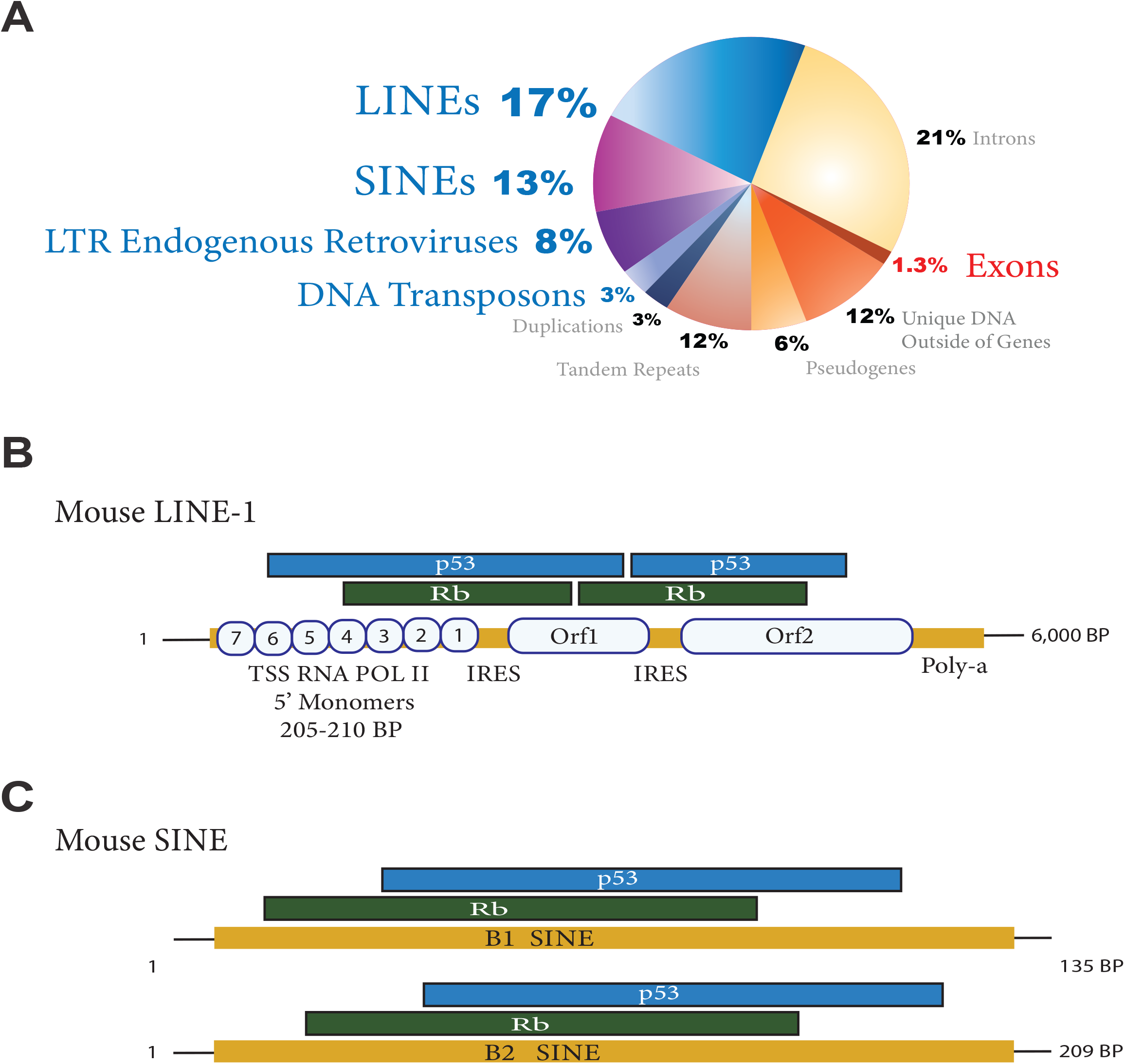
Composition of DNA in the Human Genome. **(A)** The human genome is highly repetitive with over half of the DNA consisting of repetitive elements, specifically transposons, LINEs, SINEs, and endogenous retroviruses. Retrotransposons comprise ~40% of the human genome. **(B)** Schematic of LINE1 transposable element with binding sites for the p53 and RB tumor suppressor proteins based on published ChIP data. **(C)** Schematic of B1 and B2 SINE transposable elements with binding sites for the p53 and RB tumor suppressor proteins based on published ChIP data.

Aberrant LINE1 expression occurs in approximately 50% of human cancers and correlates with loss of the tumor suppressor p53 (1). While marked derepression of TEs can be a barrier to cancer development by triggering a suicidal interferon (IFN) response (10), TEs may also promote cancer development by deregulating gene expression, deleting tumor suppressors, amplifying oncogenes, or promoting chromosomal translocations (1, 11–13). As evidence, a recent study using whole genome sequencing of human cancers implicated LINE1 derepression as a global driver of genomic instability with retrotransposition events leading to megabase-scale deletions, amplifications, and translocations across many cancer types (1). These observations present a model whereby, in addition to the canonical and rare LINE1 insertion events that have the potential to disrupt a tumor suppressor like APC (14), LINE1 derepression may promote tumorigenesis more frequently by promoting other types of genomic alterations in the cancer genome. Interestingly, chromatin immunoprecipitation (ChIP) for p53 (15) and RB((7), reanalyzed to visualize mapping to repetitive elements, shows overlapping binding of these tumor suppressors to LINE1 and SINEs (Fig.S1 Fig. 1 B&C). However, the mechanisms underlying RE derepression during cancer development and the impact of combined loss of p53 and RB remain to be fully understood.

In this study, we discover that the two most commonly mutated tumor suppressors in cancer, p53 and RB, cooperate to suppress the two most abundant classes of TE: LINE and SINEs. Further, we show that loss of p53 and RB leads to DNA hypomethylation and a decrease in the repressive histone mark, H3K9me3, which correlates with LINE1 expression. We also find that a p53 transactivation domain (TAD) mutant (p53^25,26^), which fails to increase the transcription of canonical p53 target genes after DNA damage but maintains tumor suppressor activity (16, 17), retains the ability to repress repetitive elements. These results suggest that repression of LINE1 and perhaps other repetitive elements may contribute to tumor suppression by p53 and RB.

## Results

### p53 and RB Cooperate to Suppress Expression of SINE and LINE Repetitive Elements

Expression of repetitive elements is normally suppressed in adult somatic tissues by DNA methylation and post-translational modifications of histones (18–20). Prior work has shown that 5-Azacytidine (5-Aza) treatment, a DNA demethylating agent, can induce the expression of repetitive elements in p53 deficient cells suggesting that p53 cooperates with DNA methylation machinery to silence repetitive elements (10). Specifically, 5-Aza treatment in mouse embryonic fibroblasts (MEFs) lacking p53, but not those with wild-type (WT) p53, triggered a massive induction of the SINEs B1 and B2. To confirm this observation, we generated MEFs from p53^FL/FL^ mice and infected them with an adenovirus expressing Cre recombinase (Adeno-Cre) to delete both p53 alleles. Similar to the previous report (10), Northern blot demonstrated that treatment with 10 mM 5-Aza for 48 hours induced B1 and B2 SINEs in MEFs lacking p53 but not in the parental MEFs that retained expression of p53 (Fig. 2A). Additionally, we further confirmed the previous finding (10) that cell death occurs in p53 null MEFs 120 hours after 10 mM 5-Aza (Fig. 2B), which was reported to be a consequence of activation of an endogenous type 1 interferon response.

**Figure 2.**
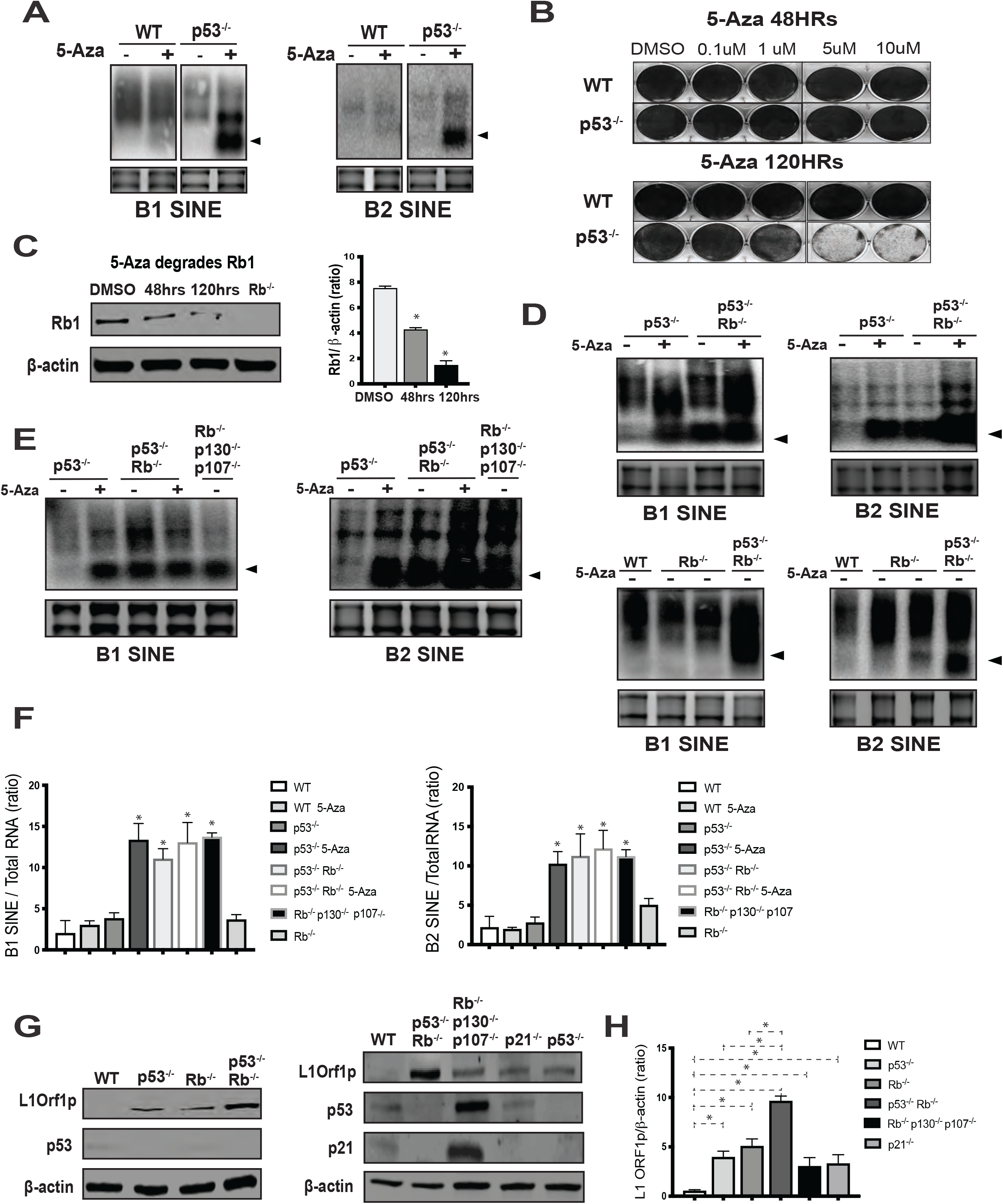
p53 and Rb cooperate to suppress B1 SINEs, B2 SINEs, and LINE1. **(A)** Transcriptional upregulation of B1 and B2 SINEs in p53 deficient MEFs treated with (+) or without (-) 10 μM 5-aza-2’-deoxycytidine (5-Aza) for 48 hours (hrs). RNA was harvested for Northern Blot with a ^32^P-labeled probe to detect B1 SINEs (left) or B2 SINEs (right). **(B)** Cytotoxicity of 5-Aza in p53 deficient MEFs was observed at 120 hrs, but not at 48 hrs. Viable cells were visualized by Coomassie blue staining. **(C)** Western blot analysis of RB and β-actin (loading control) in p53-null MEFs treated with 10 μM 5-Aza for 48 or 120 hrs. Statistical comparisons were done using the mean of each column compared to the mean of the control DMSO column analyzed via One-way ANOVA followed by Dunnett’s multiple comparisons test *(* = p < 0.05)*. **(D, E)** MEFs with the indicated genotypes were treated with (+) or without (-) 10 μM 5-Aza for 48 hours. RNA was harvested for Northern blotting using a ^32^P-labeled probe to detect B1 SINEs (left) or B2 SINEs (right). Note in MEFs lacking p53 and Rb the marked induction of B1 and B2 SINE elements even in the absence of 5-Aza. These results are representative of 3 independent experiments performed in biological replicates of n=3 p53^−/−^, n=2 p53^−/−^; Rb^−/−^ MEF lines derived from independent mouse embryos. For all Northern blots, total RNA is shown below displaying the 18S and 28S rRNA bands as loading controls stained on the gel before transfer. **(F)** Quantification of ratio of B1 or B2 SINEs to total RNA with error bar showing standard error of the mean. The mean of each column is compared to the mean of the control WT column analyzed via One-way ANOVA followed by Dunnett’s multiple comparisons test *(* = p < 0.05)*. **(G)** Western blot analysis and quantification of LINE1 ORF1p and β-actin (loading control) in MEFs with the indicated genotypes. These results are representative of 3 independent experiments performed in biological replicates of n=3 p53^−/−^, n=2 Rb^−/−^, and n=2 p53^−/−^; Rb^−/−^ MEF lines derived from independent mouse embryos. **(H)** Quantification of ratio of L1ORF1p to β-actin. Error bars represent mean ± SEM. One-way ANOVA followed by Tukey’s multiple comparisons tests for comparisons of WT vs. p53^−/−^ (P=0.003), comparison of WT vs Rb^−/−^ (P=0.0009), comparison of WT vs p53^−/−^Rb^−/−^ (P=0.0001), comparison of WT vs Rb^−/−^p130^−/−^ p107^−/−^ (P=0.015), comparison of WT vs p21^−/−^ (P=0.024), comparison of p53^−/−^ vs p53^−/−^Rb^−/−^ (P=0.0001), and comparison of Rb^−/−^ vs p53^−/−^Rb^−/−^ (P=0.0008).

Although 5-Aza may induce expression of SINEs in p53 deleted cells via DNA demethylation, other mechanisms are also possible. Notably, B1 and B2 SINE levels remain suppressed in MEFs with co-deletion of DNMT1 and p53 (21) Additionally, a previous study reported that in human cells 5-Aza reactivates gene expression at least in part via the degradation of the retinoblastoma protein RB (22). Therefore, we used Western blot to determine if 5-Aza impacts RB levels in MEFs. In p53 null MEFs, 10 mM 5-Aza decreases the levels of RB as early as 48 hours after treatment with an even more pronounced effect at 120 hours (Fig. 2C). As others had previously reported that RB recruits the histone methyltransferase EZH2 to promote histone H3 lysine 27 trimethylation (H3K27me3) to silence expression of repetitive elements (7), our results raised the possibility that p53 and RB cooperate to promote the transcriptional silencing of SINEs. Therefore, we generated MEFs from p53 ^FL/FL^; Rb^FL/FL^ mice and used Adeno-Cre to delete both alleles of each tumor suppressor gene. Remarkably, Northern blots showed that co-deletion of p53 and Rb even in the absence of 5-Aza was sufficient to induce expression of B1 and B2 SINEs (Fig. 2D). Additionally, co-deletion of all 3 Rb family members Rb1, p130, and p107 even in the presence of WT p53 and in the absence of 5-Aza was also sufficient to derepress B1 and B2 SINEs (Fig. 2E). Quantification of the ratio of B1 SINE or B2 SINE to total RNA demonstrates a statistically significant increase in expression of B1 and B2 SINEs in p53^−/−^; Rb^−/−^ MEFs compared to WT MEFs or MEFs lacking only p53 (Fig. 2F).

Having established that p53 and RB cooperate in MEFs to repress expression of B1 and B2 SINEs, we next examined whether p53 and RB cooperate to repress the expression of long interspersed nuclear elements. We used Western blot to examine the level of ORF1p as a marker for expression of full-length LINE1 elements. As previously reported (23–25), loss of p53 leads to modest increases in ORF1p expression (Fig. 2G). Similarly, we observed modest de-repression of LINE1 in cells lacking the following genes: the p53 target cyclin-dependent kinase inhibitor p21 (*Cdkn1a), Rb* alone, or the loss of all 3 *Rb* family members (*Rb, p107*, or *p130*). However, co-deletion of *p53* and *Rb* resulted in marked upregulation of ORF1p expression (Fig. 2G), which was statistically significant when compared with WT MEFs or MEFs with bi-allelic deletions of *p53* or *Rb* alone (Fig. 2H).

p53 null MEFs treated with 10 μM 5-Aza activate the expression of B1 and B2 SINEs with subsequent cell death, which has been shown to occur through a cytotoxic type 1 interferon response (10). However, here we find that expression of repetitive elements in MEFs is compatible with cell survival (Fig. S2) in the absence of 5-Aza and in the context of p53 and RB loss. Therefore, expression of repetitive elements is not sufficient to cause cell death in all contexts. Indeed, others have established that the individual loss of either RB or p53 will immortalize MEFs allowing them to grow indefinitely in tissue culture (26–28). Moreover, previous studies of MEFs lacking all three RB family members (RB, p107, and p130) demonstrated that they can form colonies in soft agar without the capacity to grow in nude mice (29). In contrast, co-deletion of p53 and RB is sufficient to fully transform MEFs because they can grow in soft agar and nude mice (30). Taken together, de-repression of repetitive elements may not be a barrier to transformation particularly when p53 and RB are co-deleted. Consistent with this notion is the observation in human cancers that the tumor suppressors *p53* and *RB* are frequently co-mutated (31–35) and that these tumors often express high levels of repetitive elements, (25, 36–38). Furthermore, because retrotransposition of LINE-1 in human cancers can lead to large chromosomal deletions, translocations, and gene amplification (1) it is conceivable that suppression of repetitive elements by p53 and RB may be an important mechanism for tumor suppression.

### p53 Transactivation Domain 1 is Required to Suppress Expression of SINEs and LINE-1 Elements

To begin to investigate whether regulation of repetitive elements may play a role in p53-mediated tumor suppression, we studied MEFs with inactivating mutations in the two N-terminal transactivation domains (TADs) of p53: p53^25,26^ and p53^53,54^. We previously showed that p53 with mutations in both TADs (p53^25,26,53,54^) was unable to transactivate known p53 targets and, like p53 deletion, lost the ability to suppress transformation in vitro and tumorigenesis in mouse models of cancer in vivo (16). However, p53 with mutation in TAD1 only (p53^25,26^), which disrupts binding to MDM2 leading to increased p53 protein expression (16, 39–42), is unable to transactivate a large majority of p53 targets in response to DNA damage but retains the ability to transactivate a small subset of p53 target genes. While p53^25,26^ is unable to induce cell cycle arrest and apoptosis in response to DNA damage (17) it nevertheless retains the ability to suppress transformation in vitro and tumorigenesis in vivo (16). Therefore, we studied whether p53^25,26^ and p53^25,26,53,54^ can suppress the expression of repetitive elements.

We generated MEFs from p53^LSL-25,26/FL^ and p53 ^LSL-25,26,53,54/FL^ embryos and infected the MEFs with Adeno-Cre in vitro to delete the p53^FL^ allele and activate the expression of the p53 TAD mutants p53^25,26^ and p53^25,26,53,54^, respectively. Western blot confirmed the expression of the p53^25,26^ and p53^25,26,53,54^ alleles in MEFs growing in tissue culture in the absence of exogenous DNA damage (Fig. 3A). Using the same cell lysates, we also examined the baseline expression level of the LINE-1 protein ORF1p. We observed that similar to wild-type p53, p53 with mutation only in TAD1 (p53^25,26^) represses ORF1p, but is unable to drive the expression of basal p21 to the same extent as wild-type p53 (Fig. 3A).In contrast, cell lysates from MEFs expressing p53 with TAD1 and TAD2 mutations (p53^25,26,53,54^) exhibited high levels of ORF1p similar to the levels in p53 null MEFs (Fig. 3A). Because TAD1 mutant p53 (p53^25,26^) represses expression of LINE-1 ORF1p and also suppresses tumorigenesis, while TAD1 and TAD2 mutant p53 (p53^25,26,53,54^) fails to repress expression of LINE-1 ORF1p or prevent tumor development, these experiments raise the possibility that p53-mediated suppression LINE-1 expression plays a critical role in p53-mediated tumor suppression.

**Figure 3.**
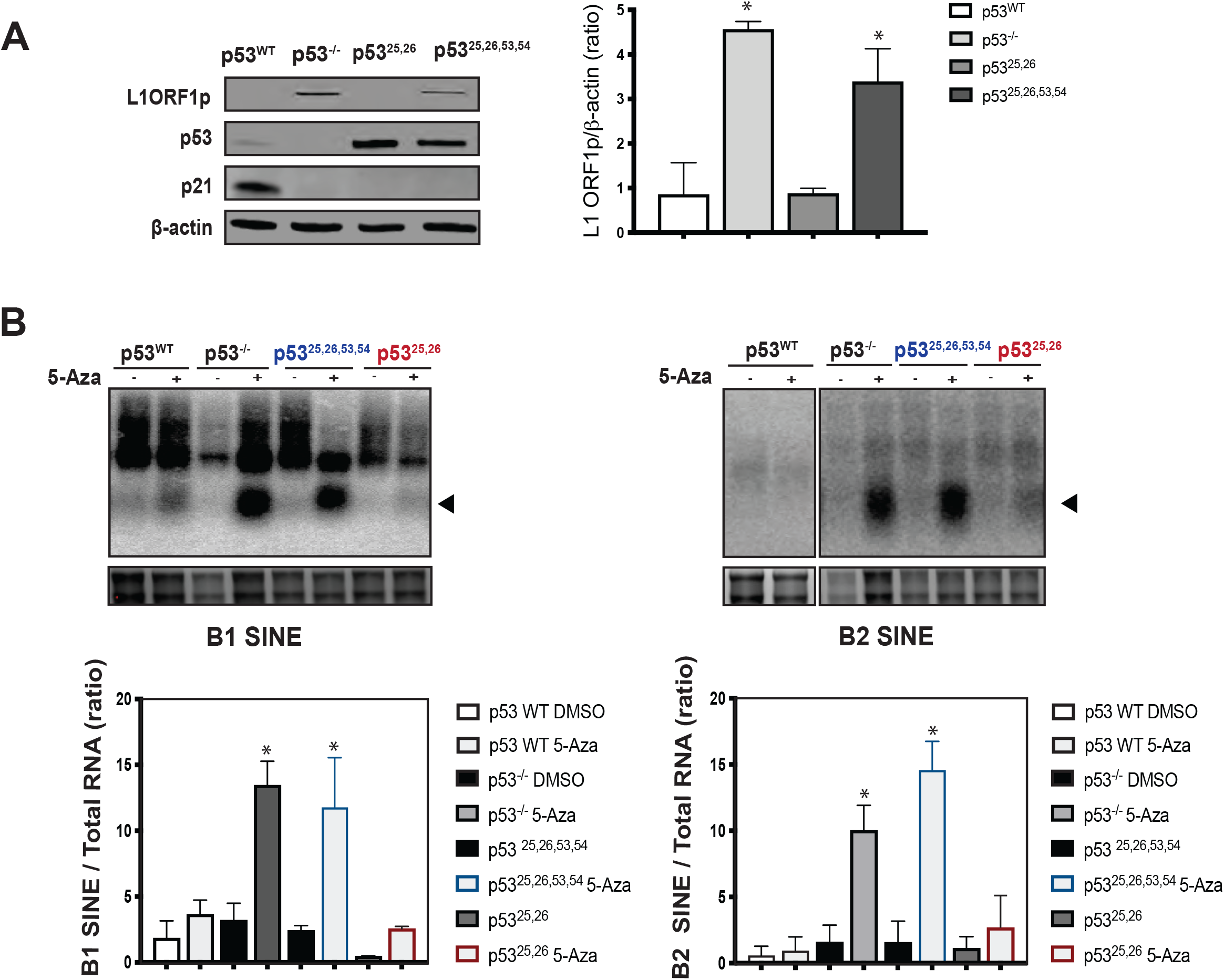
p53^25,26^, but not p53^25,26,53,54^, suppresses expression of repetitive elements. **(A)** Western blot analysis and quantification of LINE1 ORF1p and β-actin (loading control) in untreated MEFs shows that p53**^25,26^** phenocopies the ability of WT p53 to repress LINE1 ORF1p, while p53**^25,26,53,54^** modestly expresses LINE1 ORF1p, similar to p53 null. Note the stabilization of TAD p53 mutants and detection of the canonical p53 target CDKN1A (p21) in WT MEFs. This result is representative of 3 independent experiments with 2 biological replicates of p53**^25,26/-^** MEFs and 2 biological replicates of p53**^25,26,53,54/-^** MEFs. **(B)** MEFs with the indicated p53 genotype were treated with 10 μM 5-Aza for 48 hours. RNA was then harvested for Northern Blot. ^32^P-labeled probes were used to detect B1 SINEs (left) and B2 SINEs (right). Note that expression of B1 SINEs and B2 SINEs are markedly induced by 5-Aza in p53**^−/−^** MEFs and in p53**^25,26,53,54^** MEFs. However, the induction of B1 SINE and B2 SINE is much more modest in p53 WT or p53**^25,26^** MEFs. This result is representative of 3 independent experiments with 2 biological replicates of p53**^25,26/-^** MEFs and 2 biological replicates of p53**^25,26,53,54/-^** MEFs. The mean of each column is compared to the mean of the control WT column using One-way ANOVA followed by Dunnett’s multiple comparisons test *(* = p < 0.05).* Error bars represent mean ± SEM. For all Northern blots, total RNA is shown below displaying the 18S and 28S rRNA bands as loading controls stained on the gel before transfer.

We next examined the ability of p53^25,26^ and p53^25,26,53,54^ to cooperate with DNA methylation to repress SINE element expression. Intriguingly, Northern blots reveal that the p53^25,26^ mutant, when treated with 5-Aza, phenocopies the ability of WT p53 to maintain suppression of B1 or B2 expression. In contrast, the MEFs expressing the p53^25,26,53,54^ mutant exhibit a massive induction of SINEs after 5-Aza treatment, similar to MEFs lacking p53 expression (Fig 3B). Therefore, these results further extend the association between p53-mediated suppression of repetitive element expression and p53-mediated tumor suppression.

### DNA Methylation at SINE and LINE Repetitive Elements is Regulated By p53

To explore the potential role of DNA demethylation in activating expression of repetitive elements when p53 and/or RB are deleted in MEFs, we performed whole genome bisulfite sequencing (WGBS) on WT, *p53^−/−^, Rb^−/−^*, and *p53^−/−^*; *Rb^−/−^* double knockout MEFs. For WT and RB deficient MEFs, we observed high levels of DNA methylation at full length SINE B1,B2 (Fig. 4A) and LINE1 (Fig. 4B) families. In contrast, cells deficient in p53 alone or deficient in both p53 and RB demonstrated a relative decrease in DNA methylation, when compared to the WT or RB single KO (Fig. 4A and 4B). The effect of p53 deletion on DNA methylation was apparent around the promoter region of full-length LINE1 elements, however, demethylation extended across the entire 6000BP gene body (Fig. 4B). Interestingly, the loss of methylation with p53 deletion that we observed for repetitive elements holds true across the genome with loss of p53 appearing to drive a global decrease in percent methylation (Fig. S3) even in the absence of 5-Aza treatment.

**Figure 4.**
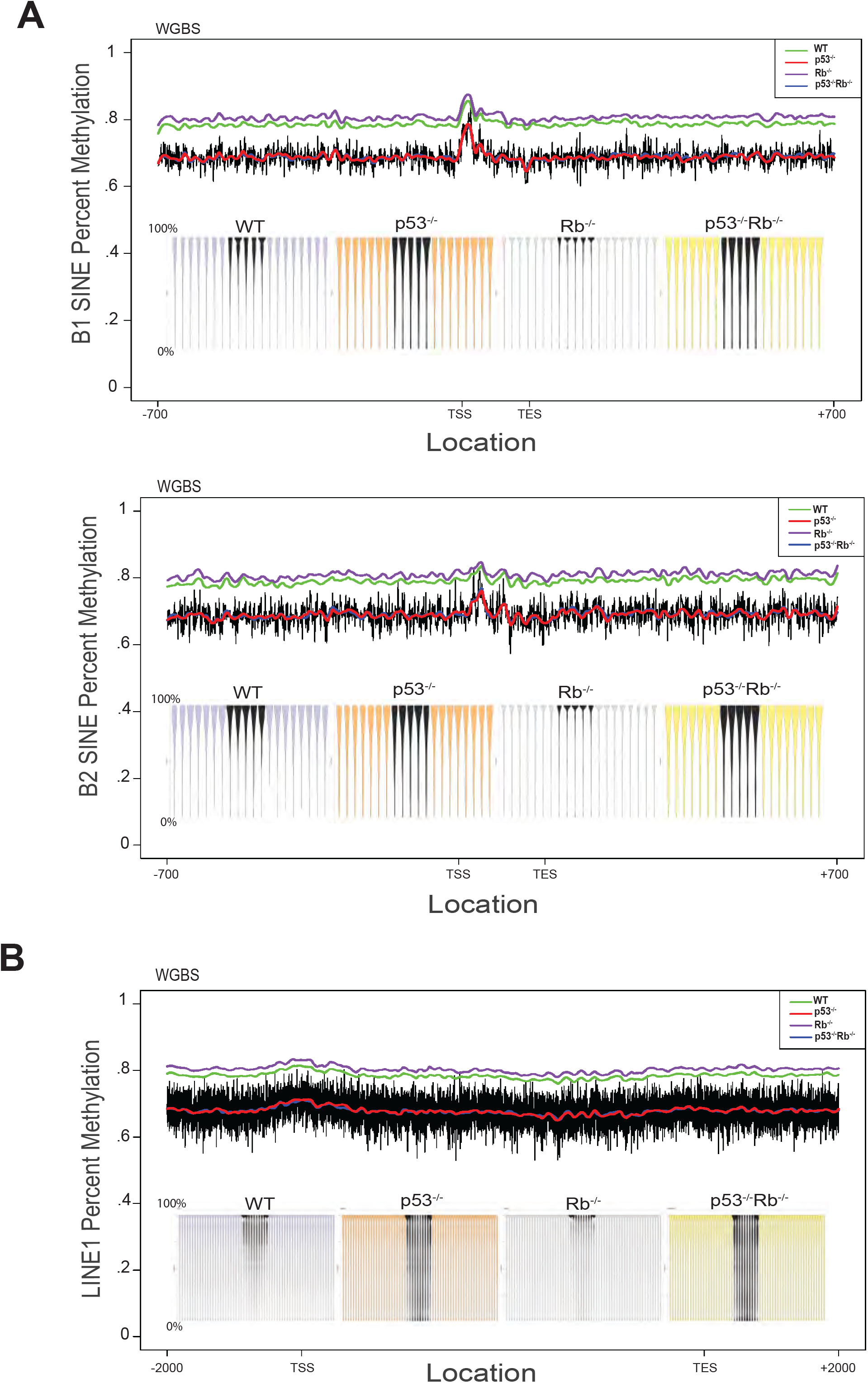
Bisulfite sequencing reveals loss of p53 causes decrease in DNA methylation at repetitive elements. DNA from the MEFs with the indicated genotypes was analyzed by WGBS. Composite analysis of all full length B1 SINE, B2 SINE, and LINE1 repetitive elements. The y-axis displays the percent methylation patterns positioned around **(A)** the B1/B2 SINE family +/-700BP and **(B)** the full length LINE1 sequence +/-2000BP. TSS (transcription start site) TES (transcription end site)Corresponding violin plots are shown as insets. Raw percent methylation values for the p53-/-sample is shown in black. The individual lines show the loess smoothed representation of percent-methylation for each genotype.

### Histone H3 Lysine 9 Methylation is Regulated by p53 and RB

Previous studies established that DNA demethylation alone is not sufficient to activate expression of B1 or B2 SINEs (21). In 2015, Varshney *et al.* used quantitative PCR (qPCR) in DNMT1 deficient MEFs to show that SINE repression did not require DNA methylation and instead SINEs are suppressed by histone methylation (21). Interestingly, they showed that DNMT1^−/−^; p53^−/−^ double knock out MEFs do not overexpress B1 or B2 SINEs(21). Because SINEs are present in the genome as hundreds of thousands of copies and are often located within introns or untranslated regions of expressed genes(43), analysis of SINE expression by qPCR is prone to error from unprocessed RNA or DNA contamination. As orthogonal validation, we used Northern and Western blots to investigate the role of histone methylation in repressing SINE and LINE1 repetitive elements in MEFs. We treated WT MEFs with 100 nM chaetocin, a specific inhibitor of the SUV39 methyltransferase family that results in selective loss of repressive H3k9me3 (44). In agreement with a previous study, chaetocin treatment led to the overexpression of B1 and B2 SINEs (Fig. 5A). Remarkably, chaetocin-induced SINE overexpression in MEFs occurred in the presence and absence of p53 and the levels of SINE expression were at least as high as the levels observed in p53^−/−^ MEFs treated with 5-Aza (Fig. 5A). Titration of chaetocin in WT MEFs shows that B1 and B2 SINEs can be derepressed to high levels with 50 nM chaetocin (Fig. 5B). Marked upregulation of LINE1 expression was also observed in WT MEFs treated with 50 nM chaetocin (Fig. 5C). Additionally, treatment with 50 nM chaetocin was toxic to WT cells (Fig. 5C bottom panel). Chaetocin toxicity was observed in WT MEFs without the 120 hour delay required to see similar effects in p53^−/−^ MEFs treated with 5-Aza (Fig. 2B). These results are consistent with a model where methylation of H3K9 and perhaps other lysine residues, exert a high degree of regulation over the repression of B1 SINE, B2 SINE and LINE1 repetitive elements.

**Figure 5.**
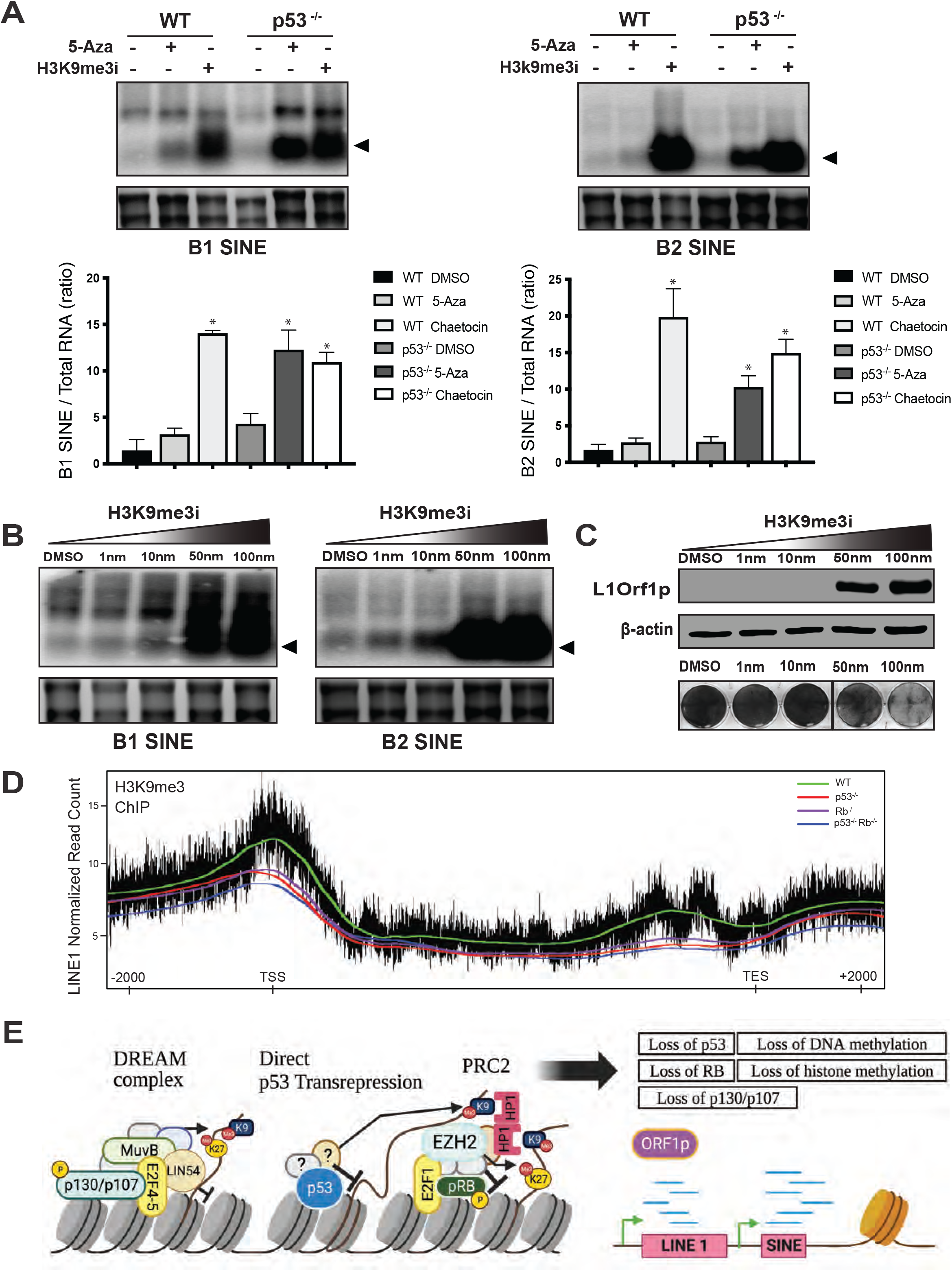
H3K9 methylation regulates expression of repetitive elements. **(A)** WT or p53^−/−^ MEFs were treated without (-) or with (+) either 10 μM 5-AZA for 48hrs or with (+) 100 nM of the H3K9me3i chaetocin for 24hrs. RNA was harvested for Northern blotting and radiolabeled using a ^32^P-labeled probe to detect B1 SINEs (left) and B2 SINEs (right). This result is representative of 3 independent experiments with 2 biological replicates and the results with 2 biological replicates per condition were quantified. Error bars represent mean ± SEM. The mean of each column is compared to the mean of the control WT column using One-way ANOVA followed by Dunnett’s multiple comparisons test *(* = p < 0.05).* **(B)** WT MEFs were treated with increasing concentrations of chaetocin for 24 hours and analyzed via Northern blot. Expression patterns of B1 SINEs (left) and B2 SINEs (right) are shown. (**C**) WT MEFs were treated for 24 hrs with increasing doses of chaetocin and analyzed via Western blot for LINE1 ORF1p and β-actin (loading control). Cytotoxicity of chaetocin was observed in WT MEFs after treatment with 50 nM chaetocin for 24hrs (bottom panel). **(D)** Chromatin immunoprecipitation of H3K9me3 followed by next generation sequencing was performed for MEFs with the indicated genotypes. Read distribution positioned +/-2000BP across full length LINE1 shows progressive loss of H3K9me3 following loss of p53 or Rb. Combined loss of p53 and Rb (blue line) shows the lowest pile up of H3K9me3. Normalized read count values for the p53-/-sample is shown in black. The individual lines show the loess smoothed representation of the normalized read count value for each genotype. (E) Working model by which p53 and RB cooperate to establish stable repression of repetitive elements. Left panel presents mechanisms by which p53 and Rb can maintain H3K9 and H3K27 trimethylation, respectively, at repetitive elements. p53 indirectly (through the DREAM complex) and/or directly (through interactions with an undiscovered cofactor) maintains H3K9me3. RB interacts with EZH2 and the polycomb repressor 2 complex (PRC2) to deposit H3K27me3. Subsequently, H3K27me3 and H3K9me3 may cooperate through HP1α dimers that maintain repressive marks and allow for the cooperative spreading of heterochromatin. Right panel shows open chromatin expressing repetitive elements following loss of tumor suppressors.

To determine if loss of the tumor suppressor genes *p53* and/or *Rb* impact global H3K9 methylation at LINE1 loci, we performed chromatin immunoprecipitation (ChIP) with an antibody to H3K9me3 using MEFs with the following genotypes: WT, *p53^−/−^, Rb^−/−^*, or double knockout *p53^−/−^*; *Rb^−/−^*. We observed that for full length LINE1 there is a stepwise depletion in the amount of H3K9me3 mapping to LINE1 as p53, RB and then both p53 and RB are deleted (Fig 5D). The depletion of H3K9me3 at full length LINE1 elements, occurs with loss of p53, RB, or both tumor suppressors and coincides with the derepression of LINE1’s ORF1p protein expression by Western blot (Fig. 1G). We observed that depletion of methylation at H3K9 extended throughout the LINE1 gene body with the most pronounced effect near the transcription start site (TSS) and promoter of LINE1.

## Discussion

Our study provides insight into the role of p53 and RB in suppressing expression of two major components of mammalian genomes: LINE1 and SINEs. Using whole genome bisulfite sequencing and H3K9me3 ChIP we show that loss of p53 in MEFs decreases DNA methylation and H3K9 methylation, which correlates with modest expression of LINE1 in the absence of 5-Aza. Remarkably, in MEFs lacking both p53 and RB, we observed robust LINE1 expression, which correlated with even greater loss of H3K9me3.

Because the histone methyltransferase chaetocin is sufficient to derepress LINE1 expression, we propose a model (Fig. 5E) where p53 and RB may restrain the expression of LINE1 and other repetitive elements at least in part through the cooperative maintenance of H3K9me3 and other repressive histone marks like H3K27me3, which have been found to be enriched at transposable elements (4, 6). In this working model, p53 may promote H3K9me3 either indirectly, by regulating the expression of key factors involved in repressive epigenetic modifying complexes like the DREAM complex (45), and/or directly through interactions with yet unknown cofactors. Furthermore, RB has already been established to direct H3K27me3 via the polycomb repressor complex 2 (PRC2) (7)(Fig. 5E). The two repressive marks could then work in tandem to contribute to the stable silencing of repetitive elements (46)(Fig.5E). Consequently, loss of either mechanism by deletion of either p53 or RB would lead to partial derepression. Consistent with this model, we find derepression of LINE1 after deletion of RB together with the two RB family members p107 and p130 that are required for the DREAM complex (45). In future studies, we will specifically test this model by deleting other components of the DREAM and PRC2 complexes.

Despite intense study for over 30 years the precise mechanisms by which p53 and RB suppress cancer remain to be fully defined. Genetically engineered mouse models of RB mutants that abrogate binding to E2F transactivation domains and are defective in transcriptional repression of E2F target genes retain tumor suppressor function(47). Similarly, p53 mutants that are not able to induce canonical transcriptional programs mediating cell cycle arrest (16, 48) apoptosis (16, 48), and senescence (48) nevertheless retain tumor suppressor function in vivo. Our data with the TAD1 p53 mutant (p53^25,26^) that loses the ability to induce cell cycle arrest and apoptosis (10, 17), but retains the ability to suppress tumorigenesis and repress LINE1 and other repetitive elements is consistent with a model where repression of LINE1 is an important mechanism of tumor suppression by p53. Overall, our results suggest that the potent tumor suppressors p53 and RB provide overlapping and independent mechanisms to repress repetitive element expression which creates redundancy and underscores the functional importance of cellular control over these elements. Indeed, our observation of marked expression of LINE1 after loss of p53 and RB, which is sufficient to transform cells and frequently occurs in human cancer, provides further impetus for future experiments to formally test whether LINE1 expression is sufficient to drive cancer development and whether regulation of LINE1 is a critical mechanism for tumor suppression by p53 and RB.

## Materials and Methods

### Cell Lines, Chemicals, and Reagents

5-aza-2’-deoxycytidine (5-aza) and chaetocin were purchased from Sigma. All drug treatments were carried out in DMEM (Invitrogen) supplemented with 10% (vol/vol) FBS and 50 μg/mL penicillin/streptomycin (Gibco). Treatment with chaetocin was for 24 hrs at 1nM to100nM as indicated in the figure legend. 5-aza was used at 10 μM for 48–120 hr, as indicated. Triple KO Rb^−/−^ p130^−/−^ p107^−/−^ MEFs were a kind gift of Julien Sage (Stanford University). All other MEFs were derived from E13.5 embryos after crossing mice with the appropriate genotype. MEFs were infected with either adeno-cre or lenti-cre to induce recombination of the floxed alleles. Where applicable, non-recombined cells were used as WT controls. Puromycin (Invitrogen) was used for the selection of cells infected with lentiviral vectors. MEFs were cultured for no more than 10 passages.

### Mice

A colony of p53 ^fl/fl^ alone, Rb^fl/fl^ alone; p53^fl/fl^; Rb^fl/fl^, p21^−/−^, p53^LSL-25,26/fl^,or p53^LSL-25,26,53,54/fl^ mice on mixed backgrounds were maintained and genotyped using Transnetyx. Animal maintenance and treatment was performed according to the protocols approved by the Institutional Animal Care and Use Committee at Duke University.

### Northern Blot Hybridization

Mouse cDNA probes for SINE B1 (sense: 5′-GCCTTTAATCCCAGCACTTG-3′, antisense: 5′-CTCTGTGTAGCCCTGGTCGT-3′), SINE B2 (sense: 5′-GCACCTGACTGCTCTTCCAGAGGT-3′, antisense: 5′-TCTTCAGACACACCAGAAGAGGGCA-3′), were generated by RT-PCR from the total RNA of Kras^G12D^; p53^−/−^ mouse sarcoma cells. The cDNAs were labeled with [α32P]-dCTP using the Random Primed DNA Labeling Kit following the protocol provided by Roche. Total RNA was TRizol (Invitrogen) extracted from MEFs with the indicated genotype after treatment with 10 μM 5-Aza for 48 hours or after treatment with an equal volume of DMSO (SIGMA). Total RNA was extracted using a Direct-zol Zymogen RNA miniprep kit (Zymo Research). Total RNA (5 μg per lane) was electrophoresed in an agarose-formaldehyde gel and transferred to a 0.45 μM Nylon membrane (Thermo Scientific). After UV crosslinking, membranes were hybridized with [α-P](Perkin Elmer)-dCTP–labeled probes and analyzed using a Typhoon Storage Phosphorimager. Quantification of all Northern blots was done using ImageJ software and reflects the relative amounts of RNA as a ratio of each band relative to the corresponding 28s and 18s loading controls.

### Cytotoxicity Assay

To determine cell cytotoxicity, cells were plated in duplicate in 6-well plates at 80% confluence and treated with increasing concentrations of drug for the durations indicated in the figure legend. At the desired time, cells were washed with PBS, fixed with 70% ethanol and stained using Coomassie Brilliant Blue (Bio-Rad). Plates were scanned in grayscale.

### Soft Agar Assay

1.8% agar (BD Diagnostics) was prepared in water. 2X cell culture medium was made by dissolving 1 bag of High glucose DMEM powder (Thermo Fisher Scientific) in half of the recommended amount of autoclaved water. The medium was sterilized using a 0.2 μm filter. Sterilized medium was mixed with 20% FBS and 1% antibiotic-antimycotic (Thermo Fisher Scientific). 2.5 ml of 0.6% agar medium consisting of 1.8% agar, 2x cell culture medium, and 1x cell culture medium was poured into 6-well plates. 0.25 ml medium containing 50,000 cells was mixed with 0.5 ml of 0.6% agar medium and was poured atop the 0.6% agar medium in the plate. Plates were incubated at 37C with 5% CO2 and were fed with 0.5 ml of 1x cell culture medium every week. Colonies in the soft agar assay were imaged after 3-4 weeks.

### Western Immunoblotting

Protein extracts were prepared by lysing cells in RIPA buffer (Sigma) supplemented with a phosSTOP tablet (Sigma-Aldrich), a protease inhibitor cocktail (Thermo FIsher Scientific) and PMSF (Sigma-Aldrich). Samples were sonicated for a total of 20 secs 10 sec on 10 sec off using a probe sonicator. Extracts were then spun down at 10,000 rpm for 10 min at 4 °C. Concentrations were determined via a BCA assay (Thermo Fisher Scientific). Equal amounts of protein were run on gradient 4–20% precast gels (Bio-Rad) and blotted/transferred to a nitrocellulose membrane (Bio-Rad). Membranes were blocked with 5% (wt/wt) BSA in Tris buffered saline and Tween 20 (TBS-T) buffer for 1 h and incubated overnight with the following primary antibodies: Anti-LINE1 ORF1p (MABC1152) (EMD Millipore), Recombinant Anti-LINE1 ORF1p antibody [EPR21844-108] (Abcam), Anti-p53NCL-p53-CM5 (Leica). Anti-Rb (IF8): sc-102 (Santa Cruz), β-Actin (8H10D10) (Cell Signaling), and p21 (12D1) (Cell Signaling). Anti-mouse and anti-rabbit secondaries were purchased from LI-COR. Blots were visualized on a LI-COR Odyssey imaging system. Quantification of all western blots was done using ImageJ software and reflects the relative amounts of protein as a ratio of each band relative to the corresponding loading control.

### ChIP assays

ChIP–seq was performed as previously described (49). Protein–DNA complexes were cross-linked using 1% formaldehyde for 10 min at room temperature and the reaction was quenched with 0.125 M glycine. Cells were washed with PBS 0.5% NP-40 and lysed by incubation in 1 M NaCl PBS/NP-40, followed by 0.1 M NaCl and 10 mM TE/NP-40. The chromatin was sheared by sonication using a Covaris focused ultrasonicator to obtain a median fragment size of 500 bp. Input samples were saved, and immunoprecipitation was performed overnight in 150 mM NaCl TE/NP-40 using 5 μg antibody bound to protein A/G sepharose. Immunoprecipitates were washed twice in RIPA buffer, twice in 250 mM LiCl with 0.5% Na-deoxycholate TE/NP-40, twice in TE and elution was performed in TE with 1% SDS. Proteinase K digestion was performed overnight at 42 °C. DNA was purified with QIAquick PCR purification kit (Qiagen). Purified DNA was sent to Genewiz for library prep and 150bp paired end sequencing.

### WGBS Library Prep and Sequencing

DNA samples were quantified using Qubit 2.0 Fluorometer (Life Technologies, Carlsbad, CA, USA). Library preparation: 100 ng gDNA was combined with 0.001 ng CpG methylated pUC19 and 0.02 ng unmethylated lambda control DNA and sheared using Covaris LE220 Focused-ultrasonicator for an average 350 bp size. The sheared material was transferred to a PCR strip tube to begin library construction. NEBNext DNA Ultra II Reagents (NEB, Ipswich, MA) were used according to the manufacturer’s instructions for end repair, A-tailing and adaptor ligation of EM-seq adaptor. The ligated samples were cleaned up according to the manufacturer’s instructions with NEBNext Sample Purification Beads. Ligated DNA was oxidized by TET2 enzymatic reaction, which was initiated by adding Fe (II) solution and then incubated for 1 h at 37 °C. Following this, Stop Reagent was added and continue incubated for 30 minutes at 37 °C. Oxidized DNA was cleaned up according to the manufacturer’s instructions with NEBNext Sample Purification Bead, denatured by Formamide, and deaminated with APOBEC enzymatic reaction, which was incubated at 37 °C for 3 hours. Deaminated DNA was cleaned up according to the manufacturer’s instructions with NEBNext Sample Purification Bead, then PCR amplified with NEBNext Q5U Master Mix and EM-Seq Index Primers. Amplified Library was cleaned up according to the manufacturer’s instructions with NEBNext Sample Purification Bead. The final library was assessed with Qubit 4.0 Fluorometer and Agilent TapeStation, and finally quantified by qPCR. Illumina sequencing: The sequencing libraries were multiplexed and sequenced on the Illumina HiSeq instrument (4000 or equivalent) according to manufacturer’s instructions. The samples were sequenced using a 2×150 Paired End (PE) configuration.

#### WGBS Data processing

Raw data was mapped to the mm10 version of the mouse genome using the BSMAP algorithm (50). Amplification artifacts were removed using the *MarkDuplicates* application from the Picard toolkit (51). The *methRatio* application from BSMAP was then used to call depth and percent methylation at each CpG loci in the mouse genome. CpG sites were filtered from subsequent analysis within a genotype if they had fewer than a depth of 4 reads in at least one replicate of that genotype.

#### ChIP-Seq Data processing

Raw H3K9me3 data was aligned to the mm10 version of the mouse genome using the Bowtie2 algorithm (52). Amplification artifacts were removed using the *MarkDuplicates* application from the Picard toolkit.Peaks were called using the *broadpeak* mode from the MACS2 algorithm (53) using default parameters. Similarly, for previously published ChIP-Seq datasets, the raw data was first downloaded from the SRA database (54). Reads were aligned to the mm10 version of the mouse genome using the Bowtie2 algorithm. Amplification artifacts were removed using the *MarkDuplicates* application from the Picard toolkit. Peaks were called using the SPP algorithm (49) using default parameters.

#### Analysis of epigenome around repetitive elements

Coordinates for B1 and B2 SINE elements as well as LINE1 elements were retrieved from the TEToolkit suite (55). Only canonical length transcripts were used for subsequent analysis. That is: 130-147nt for B1 SINEs, 150-200nt for B2 SINEs, and 5000 – 7000nt for LINE1s. For ChIP-Seq data, number of paired-end fragments with a center point at a given nucleotide relative to the repetitive element were counted. The data was then scaled by the mapped depth of the sequencing library. For WGBS data, the average percent methylation at each position was calculated. The positions upstream and downstream of the repetitive elements are calculated on a per-nucleotide basis. Within the repetitive elements, the position was scaled from 0-100% based on the location between the transcription start site (TSS) and transcription end site (TES).

#### Analysis of p53 and RB binding repetitive elements

Published ChIP seq data from WT MEFs was downloaded for p53: SRR1186260(15), Rb: SRR3175035(7), and E2F1: SRR2129988(56). Data was analyzed as described above. Bigwig files were uploaded and viewed using the UCSC genome browser. Full length LINE1 and SINE elements were manually inspected for overlapping p53 and Rb peaks.

## Data Availability

WGBS and ChIP-seq data generated in this study can be accessed at the National Center for Biotechnology Information Gene Expression Omnibus under accession no. GSEXXX (REF for the GSE deposition coming soon).

## Acknowledgements

This work was supported by grants from the National Cancer Institute 2R35 CA197616 (DGK), the National Institute of Aging 1R56 AG066487 (DGK), and the Connective Tissue Oncology Society (DGK). We thank Julien Sage for providing the Rb/p107/p130 triple knockout MEFs. Model figures were created with BioRender.com and Adobe illustrator.

## Conflict of Interests

DGK is a cofounder of and stockholder in XRAD Therapeutics, which is developing radiosensitizers. DGK is a member of the scientific advisory board and owns stock in Lumicell Inc, a company commercializing intraoperative imaging technology. None of these affiliations represents a conflict of interest with respect to the work described in this manuscript. DGK is a coinventor on a patent for a handheld imaging device and is a coinventor on a patent for radiosensitizers. XRAD Therapeutics, Merck, Bristol Myers Squibb, and Varian Medical Systems provide research support to DGK, but this did not support the research described in this manuscript. The other authors have no conflicting financial interests.

## Figure Legends

**Supplemental Figure 1.**
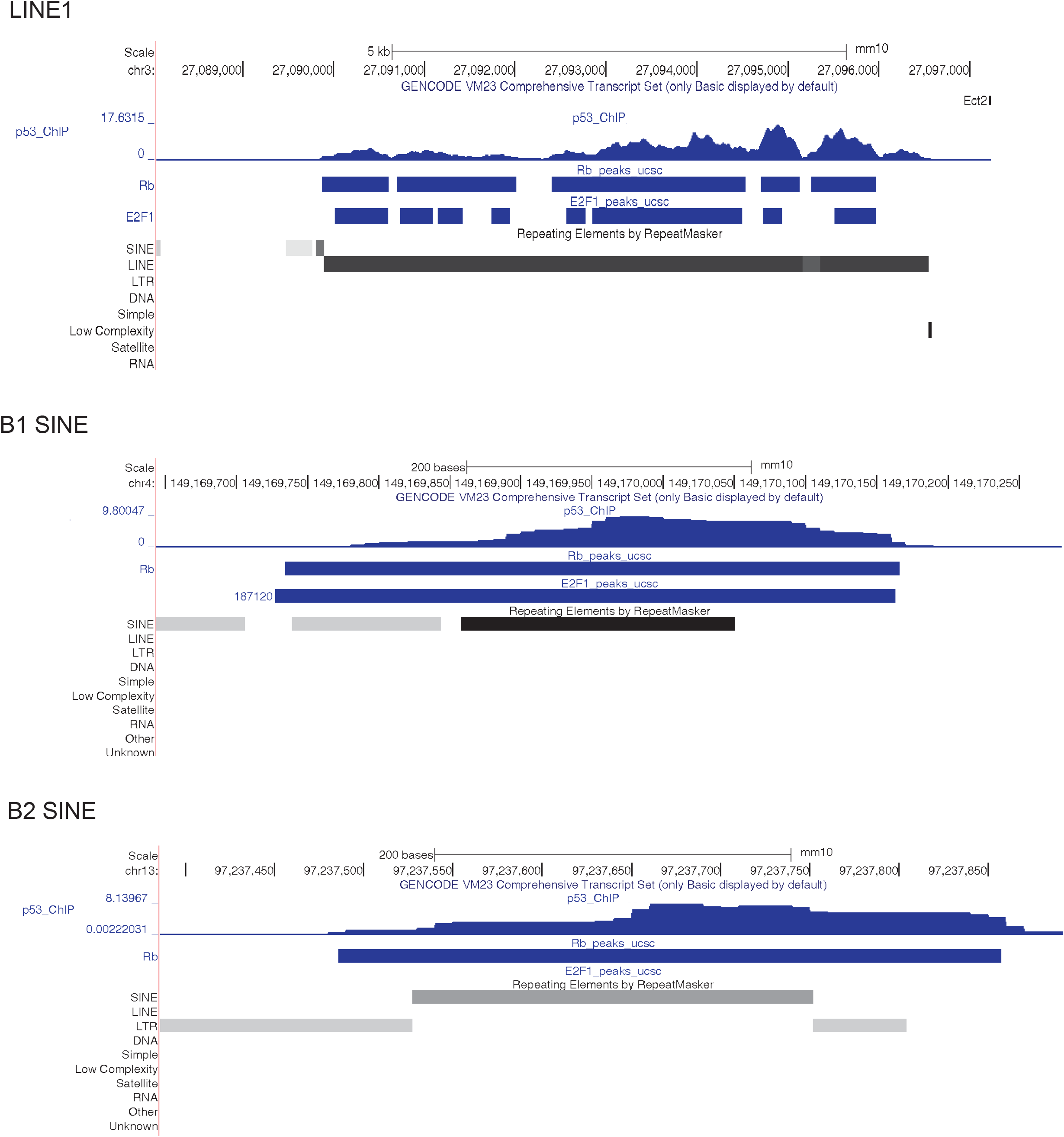
View of UCSC Genome Browser. Selected ChIP-Seq peaks show the overlap of p53, RB, and E2F1 binding sites found at LINEs and SINEs. Peaks were called using previously published data from WT MEFs. Notably RB peaks at repetitive elements do not always coincide with E2F1, B2 SINE shown as an example.

**Supplemental Figure 2.**
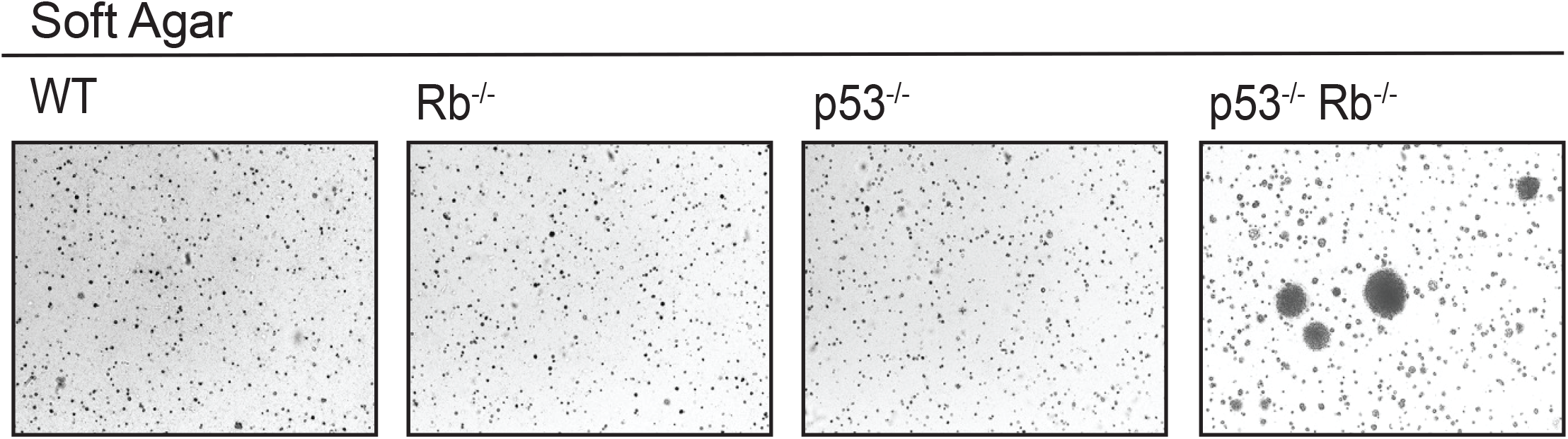
Loss of p53 and RB Transforms Cells. Soft agar assay showing transformation of MEFS following loss of both p53 and Rb. Colony formation correlates with transformation and derepression of LINE1 and SINEs.

**Supplemental Figure 3.**
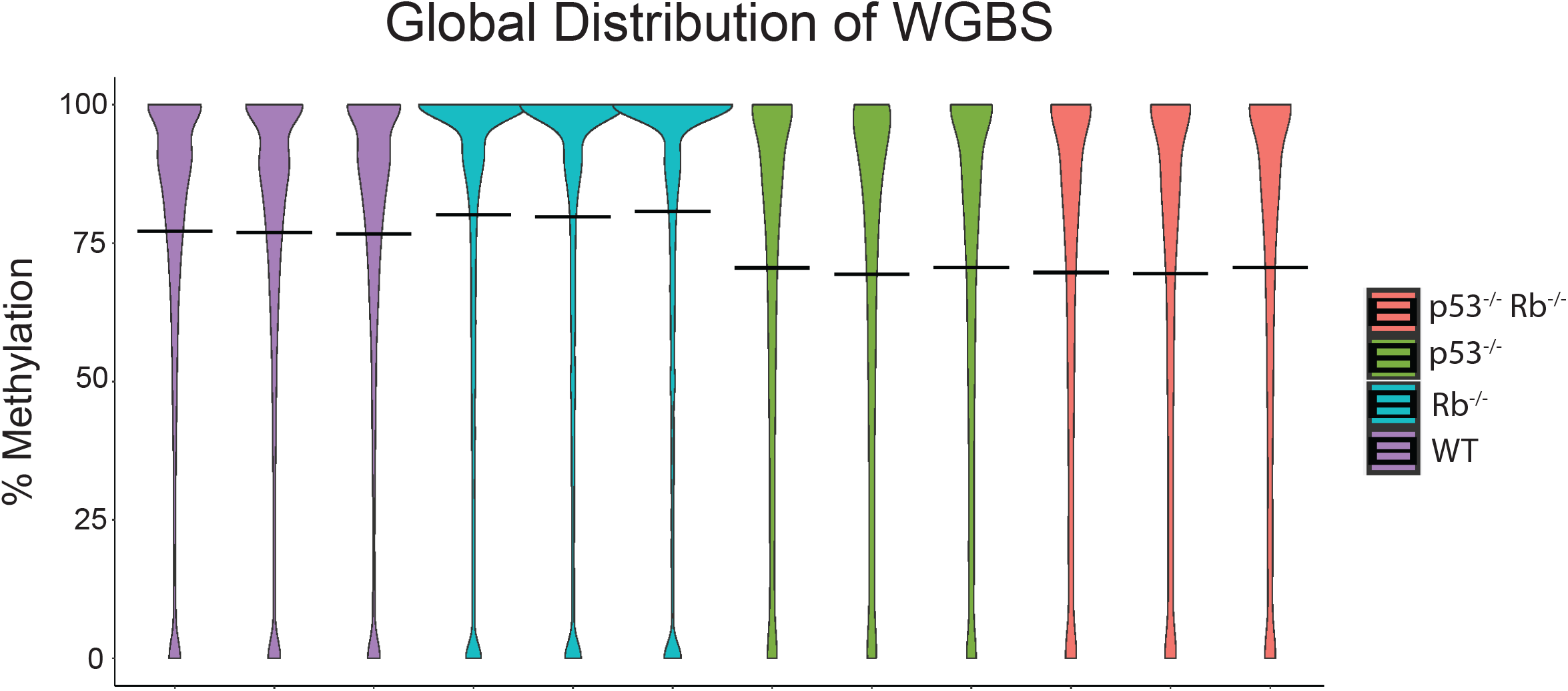
Global Distribution of WGBS in MEFs. The global distribution of the WGBS data in MEFs across genotypes. The black bar in each column marks the mean %-methylation for that sample. Loss of p53 or p53 and RB drive the largest decrease in %-methylation.

## Notes

### Competing Interest Statement

The authors have declared no competing interest.

